# Machine Learning-based Modeling of Olfactory Receptors in their Inactive State: Human OR51E2 as a Case Study

**DOI:** 10.1101/2023.02.22.529484

**Authors:** Mercedes Alfonso-Prieto, Riccardo Capelli

**Affiliations:** Computational Biomedicine, Institute for Advanced Simulation IAS-5/Institute for Neuroscience and Medicine INM-9, Forschungszentrum Jülich GmbH, Wilhelm-Johnen-Straße, D-52428 Jülich, Germany; Dipartimento di Bioscienze, Università degli Studi di Milano, Via Celoria 26, I-20133 Milan, Italy

## Abstract

Atomistic-level investigation of olfactory receptors (ORs) is a challenging task due to the experimental/computational difficulties in the structural determination/prediction for members of this family of G-protein coupled receptors. Here we have developed a protocol that performs a series of molecular dynamics simulations from a set of structures predicted *de novo* by recent machine learning algorithms and apply it to a well-studied receptor, the human OR51E2. Our study demonstrates the need for simulations to refine and validate such models. Furthermore, we demonstrate the need for the sodium ion at a binding site near D^2.50^ and E^3.39^ to stabilize the inactive state of the receptor. Considering the conservation of these two acidic residues across human ORs, we surmise this requirement also applies to the other ∼400 members of this family.

---

Olfactory receptors (ORs) are a family of G protein-coupled receptors (GPCRs) that plays a crucial role in the sense of smell. ^1^ The human genome encodes for approximately 800 GPCRs, out of which 50% are ORs.^2^ Although initially identified in the nose, ORs are expressed in different parts of the body.^3,4^ The investigation of the physiological roles of these extranasal ORs, as well as their possible involvement in pathological conditions, is attracting a growing interest.^5,6^ Moreover, given that GPCRs are the target of ∼34% of FDA-approved drugs^7^ and the wide range of biologically active molecules binding to ORs,^8^ these receptors are being explored as potential novel drug targets. ^9,10^ However, the lack of high-resolution structures for ORs has hindered the understanding of their functional mechanisms and the development of OR-targeting drugs.

Recently, the field of computational biology has made significant strides in protein structure prediction, following the development of AlphaFold2, ^11^ a deep learning (DL)-based algorithm that can predict the 3D structures of proteins from their amino acid sequences with high accuracy. The success of AlphaFold2 and other machine learning (ML)-based algorithms has provided a powerful tool to study protein structure and function. ^12–14^ Nonetheless, structural prediction of GPCRs, including ORs, still presents challenges. In particular, the algorithm predicts a single structure, despite multiple conformational states are possible for GPCRs,^15,16^ and higher average confidence scores are obtained for proteins with close homologs in the training PDB set,^17^ which is not the case for ORs.

To verify the reliability of an out-of-the-box *in silico* approach to predict OR structures and dynamics, we tested a set of models generated with six different predictors, followed by sub-microsecond molecular dynamics (MD) simulations. We chose to focus on the human olfactory receptor 51E2 (hOR51E2), associated to prostate cancer, because it has been widely studied, both experimentally and computationally. ^18,19^ Based on our test case, we propose a protocol to build reliable models of inactive, sodium-bound OR structures.

## Initial models

A set of six structural models of hOR51E2 was generated via homol-ogy modeling and ML-based prediction algorithms. For homology modeling, we relied on the SwissModel (SM) webserver,^20^ while for ML-based prediction, we considered AlphaFold (AF),^11,21^ RoseTTAFold (RF),^22^ OmegaFold (OF),^23^ and ESMFold (EF).^24^ As a last candidate, we considered a model of the receptor in its inactive state (AF_in_), generated with AlphaFold-MultiState. ^15,25^ For all the predictors considered, we tried to use the models already available to the public (*i*.*e*., without directly using the ML algorithm or modifying the default parameters – see details in the Supporting Information). In Figure 1 we show the initial predicted structures and a similarity representation among all six models, based on the calculation of the mutual backbone RMSD, followed by a 2D projection using Multidimensional Scaling (MDS).^26^ The most similar conformations are the AF and OF models (in line with OF having been trained to reproduce AF results), while the most distant ones are AF_in_ (most likely because it was trained only on inactive GPCR structures) and SM (which shows extracellular loops markedly different from the other models, inherited from the template used, see Supporting Information).

**Figure 1:**
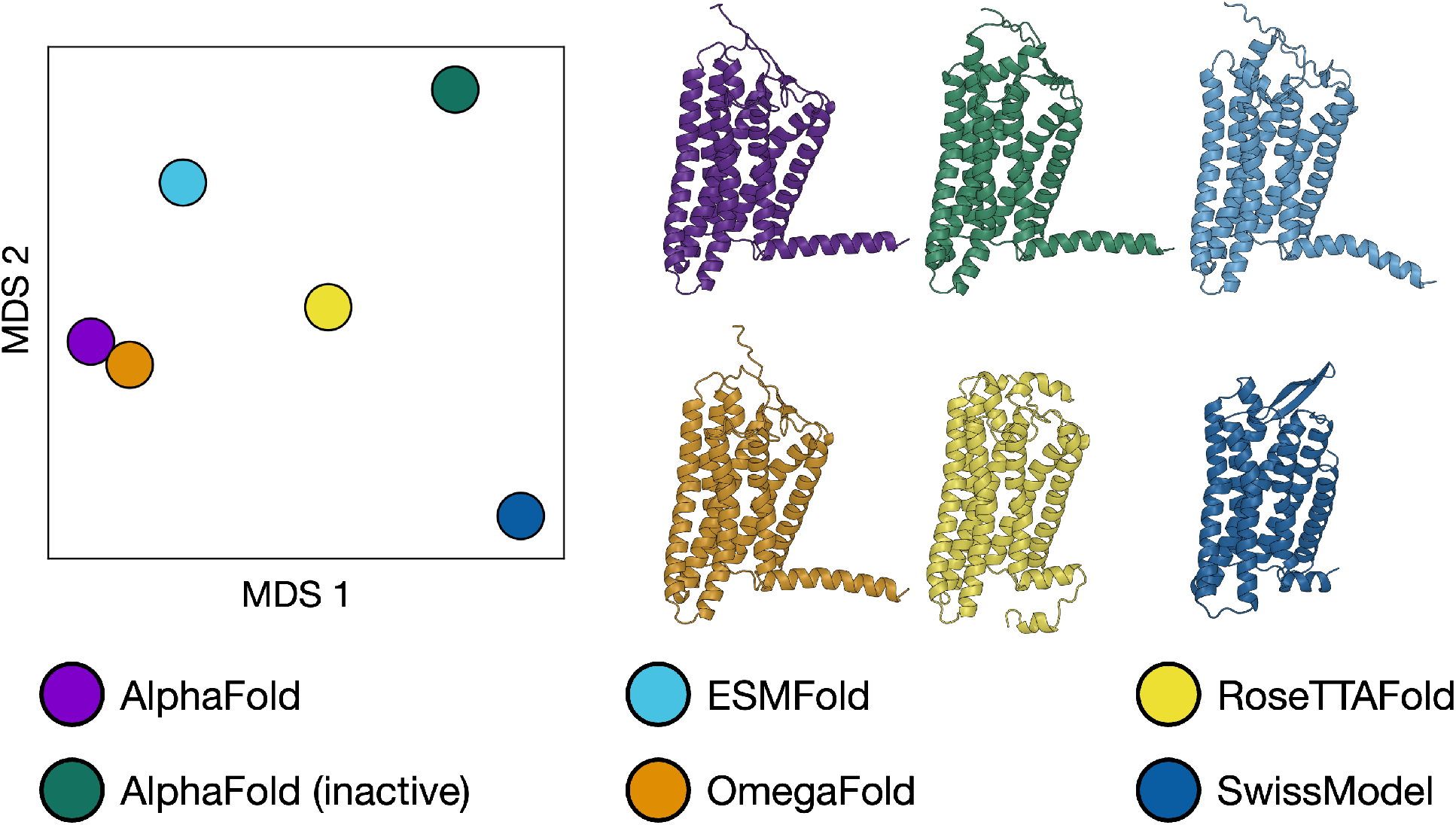
Comparison of the initial structures obtained via AI- or homology modeling-based structural prediction. *Left*, Multidimensional Scaling (MDS)-based similarity plot. *Right*, Cartoon representation of the six initial models.

### Models without Sodium ion in its Binding Site

For the first set of MD runs, we submitted the starting configurations (solvated and embedded in a POPC lipid bilayer) as set up by the CHARMM-GUI^27^ webserver (see the Methods section and the Supporting Information). During the equilibration, while the receptor and the membrane configurations were maintained in the presence of restraints, when the system was left unconstrained we observed in all cases at least a partial rearrangement of the transmembrane helices and their interfaces.

Interestingly, even before removal of the restraints on the protein structure, the interior of the receptor is flooded with water molecules passing from the intracellular to the extracellular part (Figure 2). During the last 500 ns of unconstrained simulation, the amount of flowing water increases, destabilizing the interaction network that keeps TM6 and TM7 close together, thus increasing the spacing (from 7 − 9 Å to 13 − 15 Å) between them and finally breaking the helical bundle fold.

**Figure 2:**
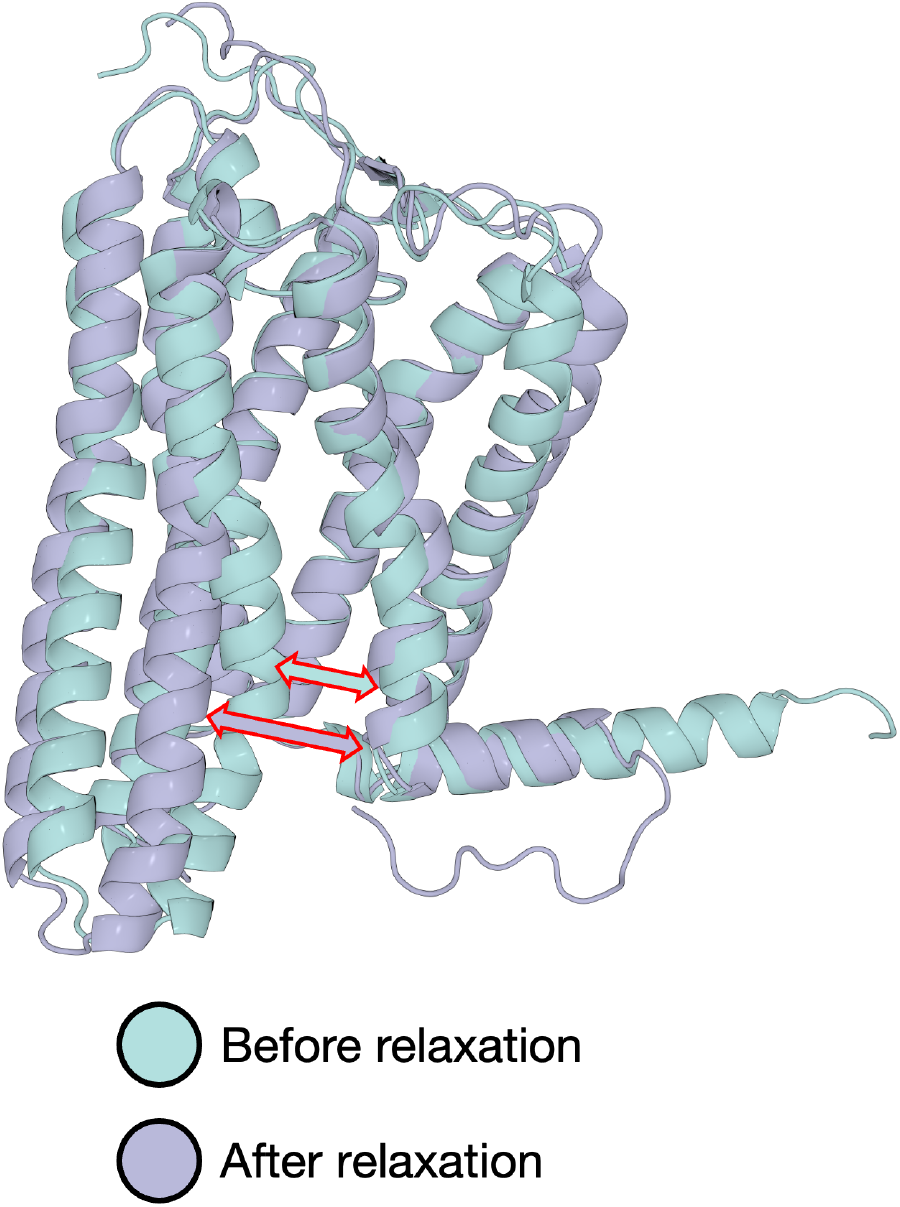
Opening of the TM6-TM7 interface in absence of sodium, exemplified here for the AF model. In all the simulations without a sodium ion bound to the receptor, the interface between TM6 and TM7 is disrupted (red-contoured arrows) and thus the receptor changes between a closed conformation (cyan) at the end of the restrained equilibration to a completely open conformation (lilac) after unrestrained MD.

One notable exception was represented by the SM structure. After ∼200 ns of unrestrained simulation, a sodium ion bound to the receptor, occupying the known ion binding pocket in class A GPCRs, close to D^2.50^ (D69 in hOR51E2). In addition, E110^3.39^ also participated in the coordination of the the Na^+^ ion. After this event, the structure appeared much more stable (despite the already broken fold). This suggests that a sodium-bound inactive structure might be more stable than an ion-devoid configuration and thus make us consider the possibility of positioning such an ion in the Na^+^ binding pocket from the beginning of the MD protocol.

### Models with Sodium in its Binding Site

In the second set of runs, we followed the same protocol but positioning a sodium ion close to the ion binding pocket in the vicinity of D69^2.50^. The time evolution of all the replicas is shown in the Supporting Information, in terms of their RMSD and A^100^ values (Figures S1 and S2). In all the 18 simulations (6 systems *×* 3 replicas per system), we observed a better preservation of the initial fold, with an RMSD of all heavy atoms around 5 Å (see Figure S2 in the SI), and the inactive conformation is maintained, as shown by the A^100^ descriptor^28^ (see Figure S1). Despite this qualitative change in the stability of the fold compared to the simulations without bound Na^+^, the sodium-bound simulations started from EF, RF, and SM configurations still showed, in all replicas, water passing from the intracellular to the extracellular part through the receptor (see Table S2 in the Supporting Information), resulting in disruption of the interface between the transmembrane helices, mainly stabilized by hydrophobic interactions.

Considering the OF and AF models, water did not pass from the intracellular part to the transmembrane part of the receptor in one and two replicas out of three, respectively, maintaining the initial fold and the TM6-TM7 distance through the whole 500 ns simulations. Finally, for AF_in_ all the 3 runs maintained the original configuration (see Table S2).

To highlight differences and similarities in the fold suggested by different structure predictors, we performed a cluster analysis of the simulations. In particular, we concatenated all the MD trajectories and calculated the reciprocal RMSD of all the frames (Figure 3), considering the heavy atoms of the transmembrane helices only and ignoring the extra- and intracellular loops, which are less stable and usually predicted with a smaller confidence.^29–31^ The results of the clustering are shown in Figure 3 and further details can be found in the Methods section. From the cluster analysis we can make two observations: (i) the three most stable models –AF_in_, AF, and OF– belong to two different clusters (1 and 4 in Figure 3); (ii) the histogram shows no overlap between the different source structures (with the exception of cluster 1, where part of RF and whole AF and OF trajectories are classified together). Therefore, upon refinement with MD, the different structure prediction methods return significantly different conformations in the transmembrane part of the receptor, even though the helical bundle should be less prone to errors in the structure prediction (and thus more stable). Interestingly, RF (which unfolds during the simulation) overlaps, at least in part, with the conformations sampled in the AF and OF simulations (see cluster 1 in Figure 3). In general, AF and OF seem to generate similar initial and MD-refined structures, that, considered together, are stable in three out of six simulations.

**Figure 3:**
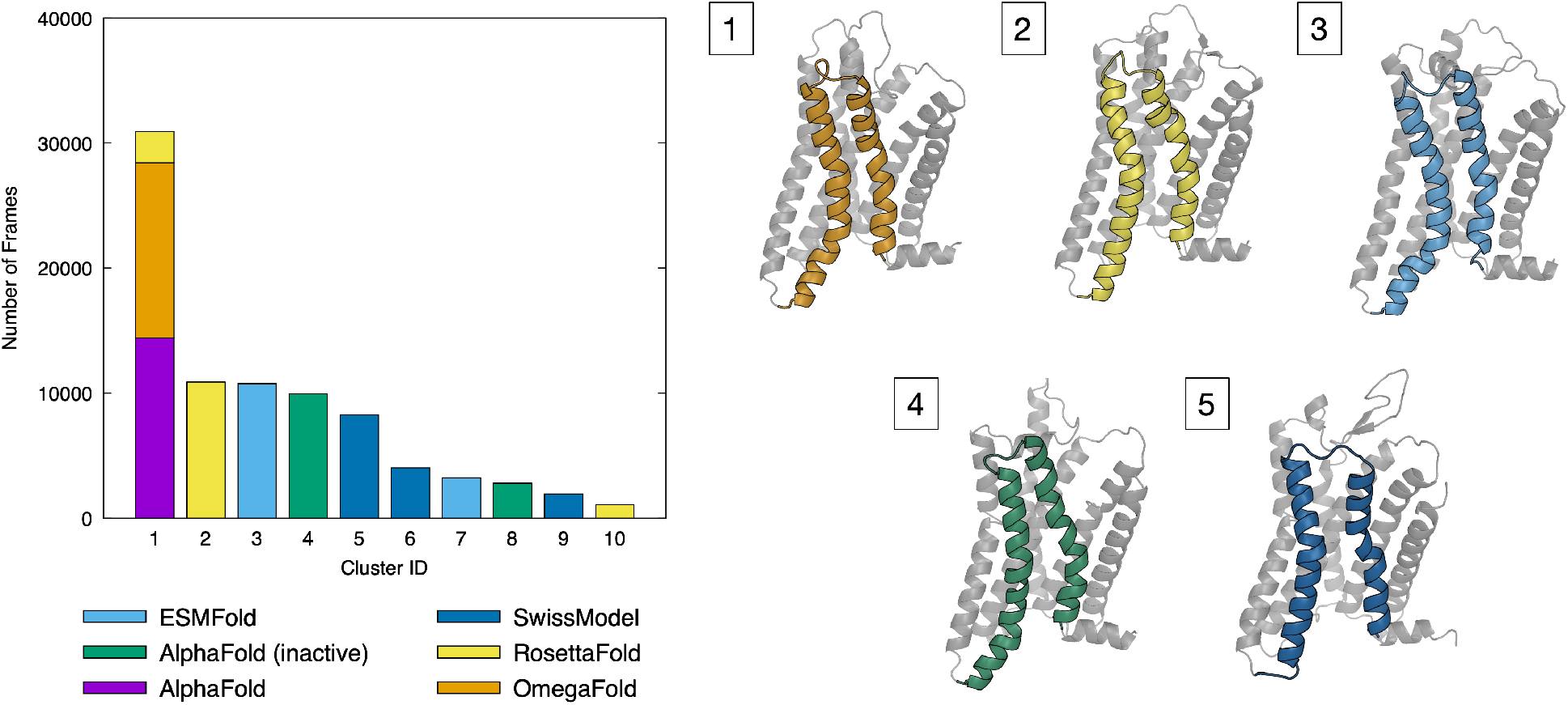
Cluster analysis for the sodium bound simulations started from different initial receptor configurations. *Left*, Cluster population; *Right*, Representative structures of clusters 1-5, with TM6 and TM7 helices colored according to the starting model.

The most evident change between the three best candidates, AF_in_ and AF/OF, is the different structural alignment of the TM6-TM7 interface, as shown by the corresponding representative structures in Figure 3 (panels 1 and 2). Contact map analysis of the centroid structures of clusters 1 and 4 (using MAPIYA^32^ (https://mapiya.lcbio.pl)) reveals a shift in the non-bonded (mainly hydrophobic) interactions that stabilize the TM6-TM7 interface in the two structures (see Figure 4). The TM6 sequence is half helical turn behind in the AF_in_ model with respect to AF/OF, whereas the TM7 helix is similar in both models. As a result, a mismatch between opposing amino acid pairs occurs at the TM6-TM7 interface. In partic-ular, the AF_in_ and AF/OF structures have almost the same TM7 residues involved in the interhelical contacts (Y279^7.40^, I286^7.48^, I290^7.52^), while for TM6 the only residue identified as interacting in both models is V246^6.43^. From a practical point of view, TM6 appears to be shifted by 1-2 residues in the structural alignment of the two models, similarly to what was observed in a recent work on another chemosensory receptor, TAS2R14,^33^ when comparing two models built with AlphaFold and I-TASSER, respectively. ^34^

**Figure 4:**
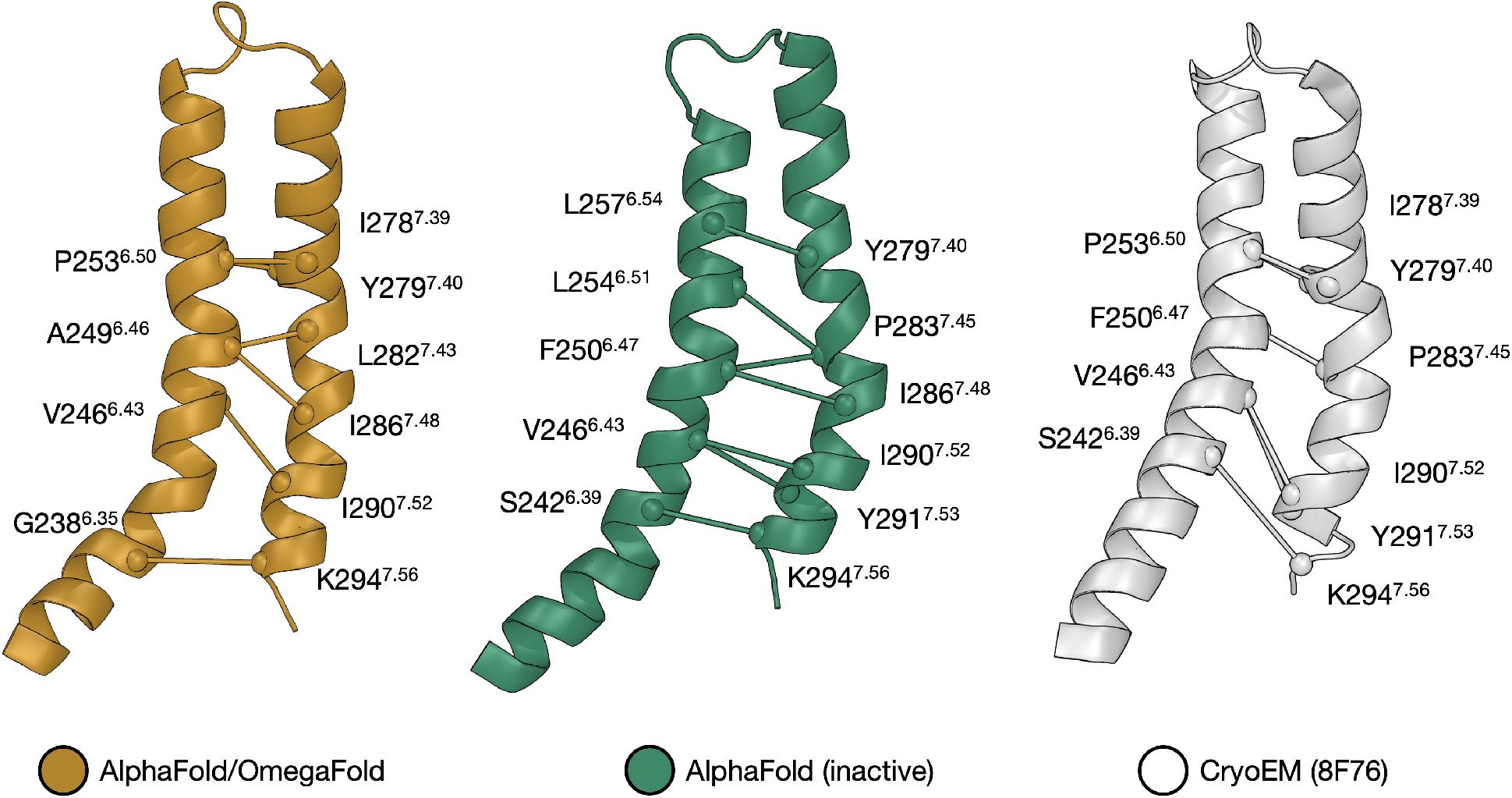
Comparison of the TM6-TM7 interface in the cryoEM structure of hOR51E2 (PDB 8F76) and the OF/AF and AF_in_ models.

Lastly, we compared the TM6-TM7 interfaces obtained for our two best *in silico* models (AF_in_ and AF/OF) with the one observed in the only experimental structure of hOR51E2 available as of April 2023 (PDB 8F76). ^19^ Taking into account that a direct comparison is not straightforward due to their different conformational state (the experimental structure corresponds to a receptor in its active state, whereas our models are in the inactive state), we can observe that the TM6-TM7 interhelical contacts are almost identical to the ones observed with AF_in_. Namely, the TM6 residues commonly identified as part of the interface for both the AF_in_ and experimental structures are S242^6.39^, V246^6.43^, and F250^6.47^. Afterwards, we observe a bending in the TM6-TM7 relative orientation (most likely related with the active state of the receptor in the cryo-EM structure); as a result P235^6.50^ becomes the last main actor of the hydrophobic interface, similarly to the AF/OF models (which are trained on both active and inactive GPCR conformations, unlike the state-specific AF_in_). A further indication that the TM6 structure is key for understanding the structure-function relationships of hOR51E2 can be found in the side chain orientation of the two TM6 residues (i.e. S258^6.55^ and R262^6.59^) involved in ligand binding based on the cryo-EM structure and mutagenesis data.^19^ In the AF/OF model, R262^6.59^ points towards the membrane, while S258^6.55^ is involved in an intrahelical H-bond with L254^6.54^. Instead, in the AF_in_ model both residues are pointing inside the binding cavity; thus, this model is more compatible with these two residues forming hydrogen bonds with the propionate ligand in the cryo-EM structure. ^19^ These observations exemplify that, although the global differences between structures generated with different predictors might seem minimal, small local differences can still result in significant changes and thus in misleading predictions regarding structure-function relationships.

In conclusion, we set up a protocol to equilibrate and test models of olfactory receptors in their inactive state embedded in a POPC membrane. We can highlight four main observations from the protocol: (i) as already suggested in previous works, ^35–37^ the reliability of structures should be tested *via* MD simulations: we observed that the initial conformation of the receptor changes in the first 50-100 ns of unrestrained simulations (see RMSD plots in the SI), confirming the need of relaxation times at least in this order of magnitude to verify the stability of a model; (ii) *de novo* structural determination can lead to significantly different predictions in presence of a multi-state system (see AF vs. AF_in_); (iii) the limited conservation of sequence motifs between human ORs and other class A GPCRs (see Table S3), especially for TM6, ^38^ can lead to gross errors in structure reconstruction; and (iv) for hOR51E2 in its inactive state –but this is most probably valid for a large set of ORs– the presence of sodium in its binding pocket is crucial for the stabilization of its fold. Sodium binding to hOR51E2 can be attributed to the presence of two negatively charged residues, D69^2.50^ and E110^3.39^. The first one is a known site for ion binding conserved in class A GPCRs, while the second position is usually occupied by S in non-olfactory class A GPCRs^39^ (see Table S4). Instead, 93% of human ORs contain Asp/Glu at both positions 2.50 and 3.39 (see Figure S6). Residue conservation in these sites can suggest a coevolutionary feature^40^ supporting the structural stability role of Na^+^ ion binding, as empirically observed by us. In line with this hypothesis, the presence of sodium in that position is also foreseen for hOR51E2 by the ML-based protein-ligand binding predictor, AlphaFill^41^ (see https://alphafill.eu/model?id=Q9H255). As a further indirect validation, a recent experimental work^42^ showed that mutation of E^3.39^ enhances the *in vitro* expression of ORs, further supporting the structural and functional importance of this residue. As pointed out by a recent commentary,^43^ *de novo* structure determination is dramatically limited by the “single answer problem”: predictors return a single structure that is, following the training, the most probable candidate. From a general point of view, this can be correct only for single-state proteins, while here (and in the majority of the biologically-relevant cases) our target GPCR has a set of different conformational states. This problem can be solved (or attenuated) taking particular care of the structural knowledge that the algorithm employs to perform its prediction. In the Heo and Feig^15^ or del Alamo *et al*.^16^ approaches, this is accomplished by limiting the training set to a single state (here GPCR experimental structures annotated to be in the inactive state), to maximize the chances of a correct prediction. The majority of the *de novo* structure determination algorithms need a properly aligned multiple sequence alignment (MSA). Hence, the abundance of sequences that can be employed in the generation of reliable MSA is a key point in the success of the structural prediction algorithms presented here. In the case of the GPCR superfamily, their predominance in the human genome (approx. 800 genes)^2^ and the availability of experimental structures (159 unique receptors in the GPCRdb, as of December 2022) provides a wealth of data to train ML predictors. However, as pointed out in references,^15,16^ GPCRs have multiple conformational states and thus special care needs to be taken when generating the corresponding MSAs. Here we further highlight that, for less similar GPCR families, such as hORs, ^44^ the limited conservation of functional motifs (or lack thereof, see Table S3) further impacts the reliability of the structural predictions. In particular, ORs lack the “rotamer toggle switch” involving W^6.48^ present on helix TM6 in non-olfactory class A GPCRs,^38,45^ but contain Y/F at positions 6.48 and 6.47 (see Table S2). Such divergence (and possible consequent MSA mismatch) may result in different structural predictions. Some of these models seem to be not good enough, as evidenced by the stability (or lack thereof) of the predicted fold of the system in MD simulations. One possible way to overcome this problem can be represented by the use of manually-curated MSA based also on structural information and/or in the training of ML weights to target specific GPCRs subfamilies in their structure predictions.

Finally, we expect that the MD-based protocol presented here for the inactive models of hOR51E2 as test case can be generalized and applied to the other ∼400 hORs, as well as to class A GPCRs. In particular, the observation that sodium binding helps stabilize the inactive models is likely to hold for the 93% of hORs with D/E at positions 2.50 and 3.39 (Figure S6) and for the 94% class A GPCRs with a negatively charged residue at position 2.50.^39^

## Methods

### System preparation

The set of six ML- and homology-based structural models of hOR51E2 generated in this work (see Supporting Information, section ‘Initial structures generation’) was preprocessed using the Protein Preparation Wizard implemented in Schrödinger Maestro 2022-3, ^46^ which automatically assigns the amino acid protonation states. Two exceptions were represented by D69^2.50^ and E110^3.39^, that were kept in their charged state. All the structures prepared were further processed via the interface of CHARMM-GUI.^27,47^ First, we built a disulfide bond between C96^3.25^ and C178^45.50^, then we defined a cubic box with dimensions 100*×*100*×* 120Å^3^, with the receptor embedded in a POPC lipid bilayer. The membrane and the receptor were solvated in water with a NaCl concentration of 0.15 M, in line with standard experimental and physiological conditions for GPCRs. The protein, lipids, and ions were parameterized using the CHARMM36m force field,^48^ while water was described with the TIP3P^49^ model.

### Molecular dynamics simulations

The simulations performed here were based on an extended version of the standard CHARMM-GUI workflow (see Supporting Information). The production step was a 500 ns-long unrestrained MD simulation with a time step of 2 fs. Velocity rescale thermostat ^50^ and cell-rescale barostat^51^ were applied to keep the temperature and pressure to 310 K and 1 bar, respectively. For all Na^+^-bound systems, we performed three independent replicas for each model, assigning different starting initial velocities. The number and simulation length of the replicas performed here is the same as recommended in the protocol used in the GPCRmd repository. ^52^ All simulations were performed using GROMACS^53^ 2021.2 patched with PLUMED. ^54,55^

### Cluster analysis

We concatenated the trajectories for all the systems simulated and performed a mutual RMSD calculation using non-hydrogen atoms of the transmembrane part of the receptor. Clustering was performed with the gromos method, ^56^ as implemented in GROMACS, using an RMSD cutoff of 2.5 Å.

## Data Availability

Data needed to reproduce the results shown in this paper (structures, topology, GROMACS and PLUMED input files, a tcl script for 7×7 RMSD calculations, etc.) and the resulting trajectories are available at Zenodo (https://doi.org/10.5281/zenodo.7817679).

## Acknowledgement

RC thanks Carlo Camilloni, Federico Ballabio, and Amit Kumawat for useful discussions. MA-P acknowledges financial support in part from the DFG Research Unit FOR2518 “Functional Dynamics of Ion Channels and Transporters – DynIon” Project P6.

## Supporting Information Available

Extensive details on the initial structures, simulation protocol steps, description of the A^100^ index for hOR51E2 and its implementation in PLUMED, figures with the time evolution of A^100^ and RMSD, a table with the water passage results, a plot that shows the amino acid conservation for positions 2.50 and 3.39, as well as tables with the Ballesteros-Weinstein numbering for hOR51E2, as listed in the GPCRdb,^25^ conserved motifs in hORs and class A GPCRs; a structural alignment of the models obtained; a MDS plot based on the RMSD of the MD trajectories; and a 7×7 RMSD matrix between the experimental cryoEM structure of hOR51E2 and the two best models obtained here (AF/OF and AF_in_).

## TOC Graphic

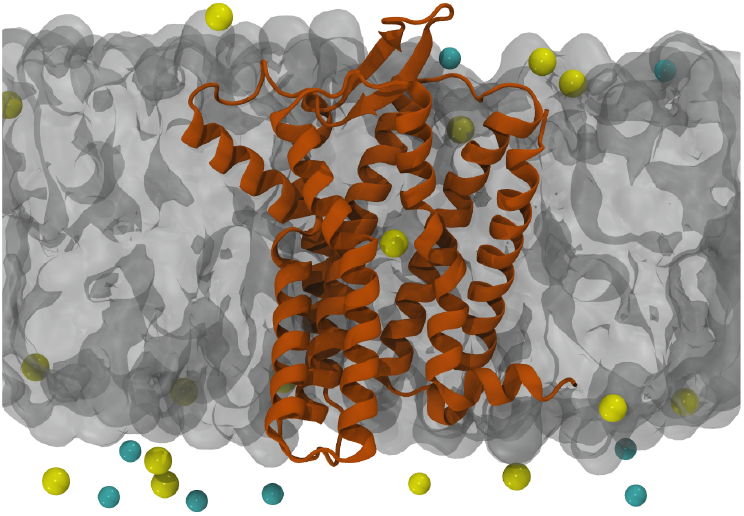

## Notes

https://doi.org/10.5281/zenodo.7817679

### Competing Interest Statement

The authors have declared no competing interest.

### Summary of Updates

New Figure 1, extensive comparison with recently published experimental structure of hOR51E2, Supporting Information file updated.

