## Supporting information file for "Machine Learning-based Modeling of Olfactory Receptors in their Inactive State: Human OR51E2 as a Case Study"

#### Initial structures generation

The hOR51E2 sequence was taken from UniProt[1] (accession code: Q9H255, amino acids 1-320). Such sequence was employed as input for all the structure prediction algorithms listed below.

**AlphaFold (AF).** We obtained the initial coordinates (accession code AF-Q9H255-F1, v3) from the AlphaFold Protein Structure Database[2] (https://alphafold.ebi.ac.uk); such model was generated with the vanilla version of Alphafold.[3]

**AlphaFold, inactive state (AF$_{\mathbf{in}}$).** AlphaFold weights were trained on all experimental structures available at the PDB, regardless of their conformation in the case of multi-state proteins. Heo and Feig recently proposed[4] to perform structure predictions considering multiple sequence alignments built on state-annotated data (*i.e.,* using only active or inactive conformations for GPCRs). Since our intention was to model the apo receptor (*i.e.,* without ligands), we therefore chose the inactive conformation. In practice, such model was obtained from GPCRdb[5] (https://gpcrdb.org), which maintains an updated database of GPCR models generated with AlphaFold-MultiState.

**RoseTTAFold (RF).** The structure predicted by RoseTTAFold[6] was obtained via the web server of the Baker Lab, ROBETTA (https://robetta.bakerlab.org).

**OmegaFold (OF).** The OmegaFold[7] prediction was run on local resources using the public implementation available on GitHub (https://github.com/HeliXonProtein/OmegaFold), release v.1.1.0, with default parameters.

**ESMFold (EF).** The structure from ESMFold[8] was obtained from the python notebook available through the ColabFold interface (https://github.com/sokrypton/ColabFold).

**SwissModel (SM).** A homology model was retrieved from the SwissModel repository.[9] The template used was the human neuropeptide Y receptor type 2 (PDB structure 7X9B, chain D), which has a sequence identity of 18.12% with hOR51E2. Unfortunately, hORs exhibit a low sequence identity ( $< 20\%$) with other class A GPCRs and thus a better template cannot be found.[10] The SM model covers residues 22-307.

### System preparation

Initially, we preprocessed all the structures obtained via *ab initio* and homology modeling algorithms by using the Protein Preparation Wizard implemented in Schrödinger Maestro version 2022-3,[11] which automatically assigns amino acid protonation states based on their microenvironment. Two exceptions were represented by D69$^{2.50}$ and E110$^{3.39}$, that were kept in the charged state. All the structures prepared in this way were further processed via the online interface of CHARMM-GUI[12,13] (https://charmm-gui.org/). First, we built a disulphide bond between C96$^{3.25}$ and C178$^{45.50}$. Then, we defined a cubic simulation box with dimensions (100 Å) × (100 Å) × (120 Å), with the receptor in the center, embedded in a POPC membrane. The lipid bilayer and the receptor were solvated in water with a NaCl concentration of 150 mM, in line with standard experimental and physiological conditions for GPCRs. The protein, lipids, and ions were parameterized using the CHARMM36m force field,[14] while water was described with the TIP3P[15] model.

#### Molecular dynamics simulations

Our protocol is a modified and extended version of the standard CHARMM-GUI-suggested workflow for equilibration and production runs of transmembrane proteins. It comprises nine different steps (here we consider the same numbering used in the CHARMM-GUI input files, 0-6, plus two additional steps 7-8):

- step 0: 5,000 steps of steepest descent minimization, restraining the protein backbone ($k = 4,000$ kJ/mol/nm$^2$) and sidechains ($k = 2,000$ kJ/mol/nm$^2$), lipid phosphate groups ($k = 1,000$ kJ/mol/nm$^2$) and dihedrals ($k = 1,000$ kJ/mol/rad$^2$).

- step 1: 125 ps of MD with a time step of 1 fs, restraining the protein backbone ($k = 4,000$ kJ/mol/nm$^2$) and sidechains ($k = 2,000$ kJ/mol/nm$^2$), lipid phosphate groups ($k = 1,000$ kJ/mol/nm$^2$) and dihedrals ($k = 1,000$ kJ/mol/rad$^2$).

- step 2: 125 ps of MD with a time step of 1 fs, restraining the protein backbone ($k = 2,000$ kJ/mol/nm$^2$) and sidechains ($k = 1,000$ kJ/mol/nm$^2$), lipid phosphate groups ($k = 400$ kJ/mol/nm$^2$) and dihedrals ($k = 400$ kJ/mol/rad$^2$).

- step 3: 125 ps of MD with a time step of 1 fs, restraining the protein backbone ($k = 1,000$ kJ/mol/nm$^2$) and sidechains ($k = 500$ kJ/mol/nm$^2$), lipid phosphate groups ($k = 400$ kJ/mol/nm$^2$) and dihedrals ($k = 200$ kJ/mol/rad$^2$).

- step 4: 500 ps of MD with a time step of 2 fs, restraining the protein backbone ($k = 500$ kJ/mol/nm$^2$) and sidechains ($k = 200$ kJ/mol/nm$^2$), lipid phosphate groups ($k = 200$

kJ/mol/nm$^2$) and dihedrals ($k = 200$ kJ/mol/rad$^2$).

- step 5: 500 ps of MD with a time step of 2 fs, restraining the protein backbone ($k = 200$ kJ/mol/nm$^2$) and sidechains ($k = 50$ kJ/mol/nm$^2$), lipid phosphate groups ($k = 40$ kJ/mol/nm$^2$) and dihedrals ($k = 100$ kJ/mol/rad$^2$).

- step 6: 100 ns of MD with a time step of 2 fs, restraining the protein backbone ($k = 50$ kJ/mol/nm$^2$); this step is 10-fold longer than in the standard CHARMM-GUI protocol.

- step 7: 100 ns of MD with a time step of 2 fs, restraining the protein backbone ($k = 5$ kJ/mol/nm$^2$); this is a completely new step, which increases the length of the restrained equilibration.

The last, production step (8) consists of $2.5 \cdot 10^8$ steps with a time step of 2 fs (500 ns) of unrestrained MD simulation. For both van der Waals and short-range interactions, a cutoff distance was set at 10 Åand long-range electrostatic interactions were computed via the smooth particle mesh Ewald summation method.[16] Temperature was kept at 310 K via velocity rescale thermostat[17] with a coupling time of 0.2 ps on three groups: protein, membrane, and water and ions. Pressure was kept constant at 1 bar with the semi-isotropic cell rescale barostat[18] with a coupling time of 0.5 ps. All the simulations were performed using GROMACS[19] version 2021.2; three replicas were run for each model.

### $A^{100}$ activation index

$A^{100}$ is an estimator of the activation state of class A GPCR structures,[20] defined as a weighted sum of distances between $C_\alpha$ atoms of selected residues, namely:

$$\begin{aligned} A^{100} = & -14.43 \cdot d(V^{1.53}, L^{7.55}) - 7.62 \cdot d(D^{2.50}, T^{3.37}) + \\ & + 9.11 \cdot d(N^{3.42}, I^{4.42}) - 6.32 \cdot d(W^{5.66}, A^{6.34}) + \\ & - 5.22 \cdot d(L^{6.58}, Y^{7.35}) + 278.88 \end{aligned} \quad (1)$$

where $d(X, Y)$ is the Euclidean distance between residues $X$ and $Y$. In equation (1) the amino acid names are based on the generic residue numbering proposed by Ballesteros and Weinstein[21] (BW) for class A GPCRs. In the case of hOR51E2, the BW numbering was taken from GPCRdb[5] (see Table S1). We would like to note here that, because of the lack of conservation of some class A GPCR motifs in ORs (see Table S3), the nature of some residues is different, yet the BW position is the same. Thus, the $A^{100}$ activation index for hOR51E2 is defined as:

$$\begin{aligned} A^{100}_{OR51E2} = & -14.43 \cdot d(V44^{1.53}, A293^{7.55}) - 7.62 \cdot d(D69^{2.50}, A108^{3.37}) + \\ & + 9.11 \cdot d(I113^{3.42}, T140^{4.42}) - 6.32 \cdot d(L225^{5.66}, F237^{6.34}) + \\ & - 5.22 \cdot d(H261^{6.58}, V274^{7.35}) + 278.88 \end{aligned} \quad (2)$$

This estimator has been implemented in PLUMED 2.8.[22,23]

As shown in Figure S1, the $A^{100}$ index remains around a value of 11 throughout the entire 500 ns simulation for all six systems, regardless of the replica considered. Following the two-state model of the original work,[20] where a class A GPCR structure is defined as in the inactive state for an index value smaller than 30, we can consider all the simulated models in the inactive state.

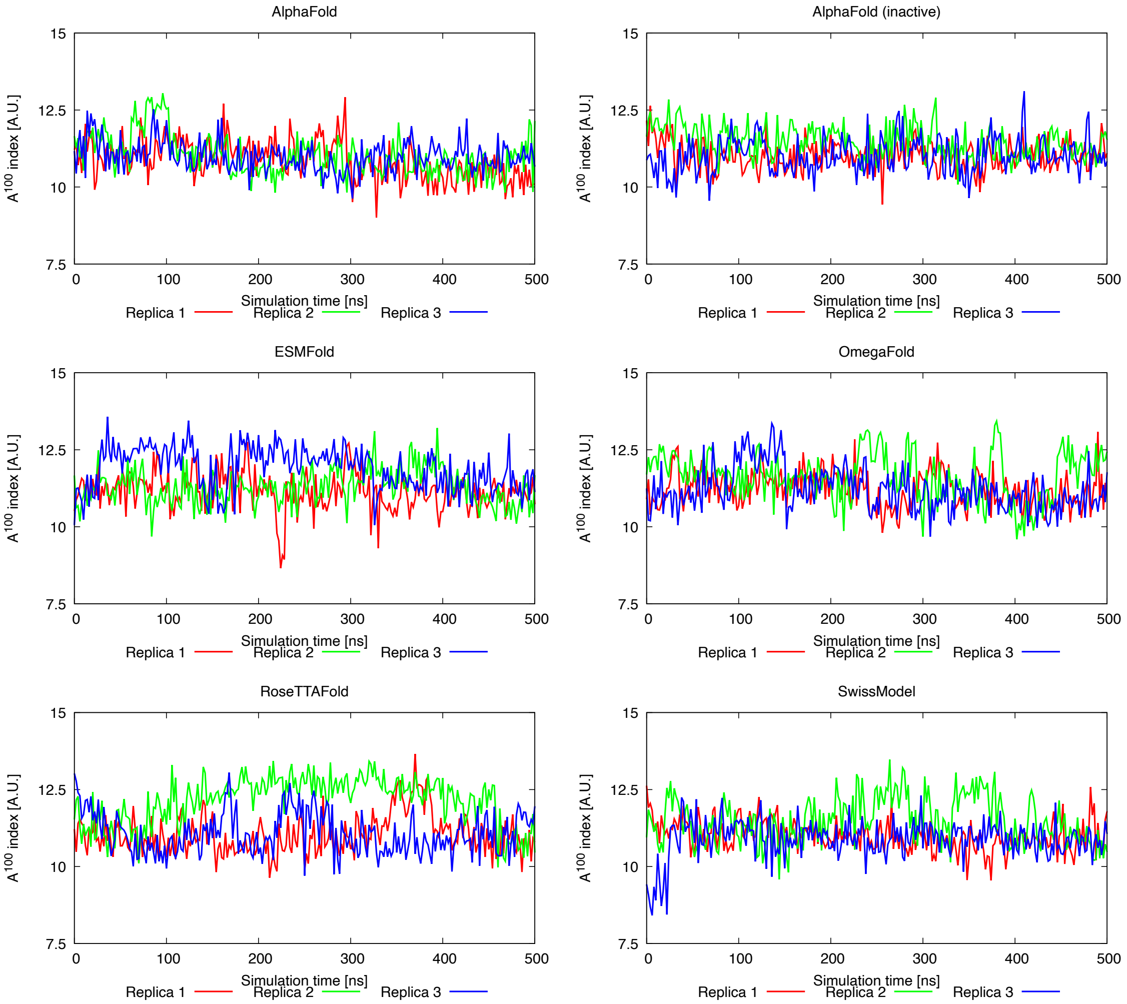


Figure S1: Time evolution of the $A^{100}$ activation index for the eighteen MD simulations of sodium-bound hOR51E2 performed in this work.

### $A^{100}$ implementation in PLUMED for hOR51E2

```
UNITS LENGTH=A

WHOLEMOLECULES ENTITY0=1-4571


# We are here considering the C-alpha distances suggested in Ibrahim et al. JCIM 2019

# the numbering refers to hOR51E2

d1: DISTANCE ATOMS=340,4325

d2: DISTANCE ATOMS=757,1381

d3: DISTANCE ATOMS=1450,1881

d4: DISTANCE ATOMS=3246,3440

d5: DISTANCE ATOMS=3801,4019


# And this is the eq.1 of the same work cited above

MATHEVAL ...

  ARG=d1,d2,d3,d4,d5

  VAR=x1,x2,x3,x4,x5

  FUNC=-14.43*x1-7.62*x2+9.11*x3-6.32*x4-5.22*x5+278.88

  LABEL=a100

  PERIODIC=NO

... MATHEVAL


PRINT FILE=a100.dat ARG=d1,d2,d3,d4,d5,a100 STRIDE=1


ENDPLUMED
```

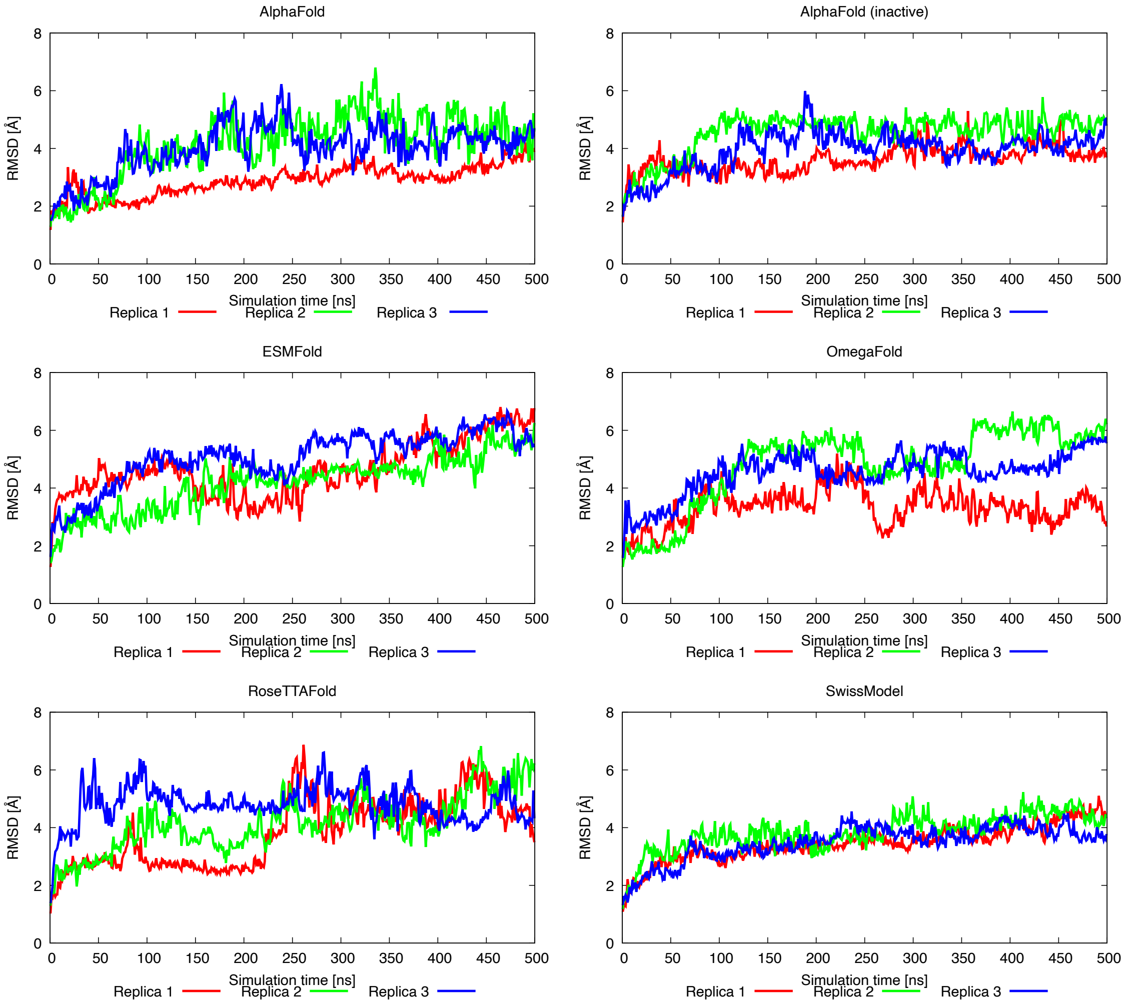


Figure S2: Time evolution of the RMSD of the heavy atoms of the whole protein for the eighteen MD simulations of sodium-bound hOR51E2 performed in this work.

Table S1: Ballesteros-Weinstein generic numbering for hOR51E2, as listed in the GPCRdb[5] (https://gpcrdb.org/residue/residuetabledisplay, accessed on March 2023).

| **TM1** | | **TM2** | | **TM3** | | **TM4** | | **TM5** | | **TM6** | | **TM7** | |
|---|---|---|---|---|---|---|---|---|---|---|---|---|---|
| 1x32 | H23 | 2x37 | A56 | 3x21 | S92 | 4x38 | N136 | 5x32 | T191 | 6x26 | S229 | 7x29 | H268 |
| 1x33 | F24 | 2x38 | P57 | 3x22 | F93 | 4x39 | N137 | 5x33 | L192 | 6x27 | K230 | 7x30 | P269 |
| 1x34 | W25 | 2x39 | M58 | 3x23 | E94 | 4x40 | T138 | 5x34 | P193 | 6x28 | S231 | 7x31 | I270 |
| 1x35 | V26 | 2x40 | Y59 | 3x24 | A95 | 4x41 | V139 | 5x35 | N194 | 6x29 | E232 | 7x32 | V271 |
| 1x36 | G27 | 2x41 | L60 | 3x25 | C96 | 4x42 | T140 | 5x36 | V195 | 6x30 | R233 | 7x33 | R272 |
| 1x37 | F28 | 2x42 | F61 | 3x26 | L97 | 4x43 | A141 | 5x37 | V196 | 6x31 | A234 | 7x34 | V273 |
| 1x38 | P29 | 2x43 | L62 | 3x27 | T98 | 4x44 | Q142 | 5x38 | Y197 | 6x32 | K235 | 7x35 | V274 |
| 1x39 | L30 | 2x44 | C63 | 3x28 | Q99 | 4x45 | I143 | 5x39 | G198 | 6x33 | A236 | 7x36 | M275 |
| 1x40 | L31 | 2x45 | M64 | 3x29 | M100 | 4x46 | G144 | 5x40 | L199 | 6x34 | F237 | 7x37 | G276 |
| 1x41 | S32 | 2x46 | L65 | 3x30 | F101 | 4x47 | I145 | 5x41 | T200 | 6x35 | G238 | 7x38 | D277 |
| 1x42 | M33 | 2x47 | A66 | 3x31 | F102 | 4x48 | V146 | 5x42 | A201 | 6x36 | T239 | 7x39 | I278 |
| 1x43 | Y34 | 2x48 | A67 | 3x32 | I103 | 4x49 | A147 | 5x43 | I202 | 6x37 | C240 | 7x40 | Y279 |
| 1x44 | V35 | 2x49 | I68 | 3x33 | H104 | 4x50 | V148 | 5x44 | L203 | 6x38 | V241 | 7x41 | L280 |
| 1x45 | V36 | 2x50 | D69 | 3x34 | A105 | 4x51 | V149 | 5x45 | L204 | 6x39 | S242 | 7x42 | L281 |
| 1x46 | A37 | 2x51 | L70 | 3x35 | L106 | 4x52 | R150 | 5x46 | V205 | 6x40 | H243 | 7x43 | L282 |
| 1x47 | M38 | 2x52 | A71 | 3x36 | S107 | 4x53 | G151 | 5x47 | M206 | 6x41 | I244 | 7x45 | P283 |
| 1x48 | F39 | 2x53 | L72 | 3x37 | A108 | 4x54 | S152 | 5x48 | G207 | 6x42 | G245 | 7x46 | P284 |
| 1x49 | G40 | 2x54 | S73 | 3x38 | I109 | 4x55 | L153 | 5x49 | V208 | 6x43 | V246 | 7x47 | V285 |
| 1x50 | N41 | 2x55 | T74 | 3x39 | E110 | 4x56 | F154 | 5x50 | D209 | 6x44 | V247 | 7x48 | I286 |
| 1x51 | C42 | 2x551 | S75 | 3x40 | S111 | 4x57 | F155 | 5x51 | V210 | 6x45 | L248 | 7x49 | N287 |
| 1x52 | I43 | 2x56 | T76 | 3x41 | T112 | 4x58 | F156 | 5x52 | M211 | 6x46 | A249 | 7x50 | P288 |
| 1x53 | V44 | 2x57 | M77 | 3x42 | I113 | 4x59 | P157 | 5x53 | F212 | 6x47 | F250 | 7x51 | I289 |
| 1x54 | V45 | 2x58 | P78 | 3x43 | L114 | 4x60 | L158 | 5x54 | I213 | 6x48 | Y251 | 7x52 | I290 |
| 1x55 | F46 | 2x59 | K79 | 3x44 | L115 | 4x61 | P159 | 5x55 | S214 | 6x49 | V252 | 7x53 | Y291 |
| 1x56 | I47 | 2x60 | I80 | 3x45 | A116 | 4x62 | L160 | 5x56 | L215 | 6x50 | P253 | 7x54 | G292 |
| 1x57 | V48 | 2x61 | L81 | 3x46 | M117 | 4x63 | L161 | 5x57 | S216 | 6x51 | L254 | 7x55 | A293 |
| 1x58 | R49 | 2x62 | A82 | 3x47 | A118 | 4x64 | I162 | 5x58 | Y217 | 6x52 | I255 | 7x56 | K294 |
| 1x59 | T50 | 2x63 | L83 | 3x48 | F119 | 4x65 | K163 | 5x59 | F218 | 6x53 | G256 | | |
| 1x60 | E51 | 2x64 | F84 | 3x49 | D120 | 4x66 | R164 | 5x60 | L219 | 6x54 | L257 | | |
| | | 2x65 | W85 | 3x50 | R121 | 4x67 | L165 | 5x61 | I220 | 6x55 | S258 | | |
| | | 2x66 | F86 | 3x51 | Y122 | | | 5x62 | I221 | 6x56 | V259 | | |
| | | 2x67 | D87 | 3x52 | V123 | | | 5x63 | R222 | 6x57 | V260 | | |
| | | | | 3x53 | A124 | | | 5x64 | T223 | 6x58 | H261 | | |
| | | | | 3x54 | I125 | | | 5x65 | V224 | 6x59 | R262 | | |
| | | | | 3x55 | C126 | | | 5x66 | L225 | 6x60 | F263 | | |
| | | | | 3x56 | H127 | | | 5x67 | Q226 | 6x61 | G264 | | |
| | | | | | | | | 5x68 | L227 | | | | |
| **ICL1** | | | | **ICL2** | | **ECL2** | | | | | | **H8** | |
| 12x48 | R52 | | | 34x50 | P128 | 45x50 | C178 | | | | | 8x47 | T295 |
| 12x49 | S53 | | | 34x51 | L129 | 45x51 | V179 | | | | | 8x48 | K296 |
| 12x50 | L54 | | | 34x52 | R130 | 45x52 | H180 | | | | | 8x49 | Q297 |
| 12x51 | H55 | | | 34x53 | H131 | | | | | | | 8x50 | I298 |
| | | | | 34x54 | A132 | | | | | | | 8x51 | R299 |
| | | | | 34x55 | A133 | | | | | | | 8x52 | T300 |
| | | | | 34x56 | V134 | | | | | | | 8x53 | R301 |
| | | | | 34x57 | L135 | | | | | | | 8x54 | V302 |
| | | | | | | | | | | | | 8x55 | L303 |
| | | | | | | | | | | | | 8x56 | A304 |
| | | | | | | | | | | | | 8x57 | M305 |
| | | | | | | | | | | | | 8x58 | F306 |
| | | | | | | | | | | | | 8x59 | K307 |
| | | | | | | | | | | | | 8x60 | I308 |

Table S2: Water passage observed in MD simulations.

| Predictor | Replica 1 | Replica 2 | Replica 3 |
|---|---|---|---|
| AF | Yes | **No** | **No** |
| $AF_{in}$ | **No** | **No** | **No** |
| EF | Yes | Yes | Yes |
| OF | **No** | Yes | Yes |
| RF | Yes | Yes | Yes |
| SM | Yes | Yes | Yes |

#### Comparison of the *in silico* models with sodium ion bound

We assessed the similarity of the conformations explored during the MD simulations by calculating the mutual backbone RMSD and using Multidimensional Scaling (MDS), a non-linear dimensionality reduction approach to visualize the information contained in the RMSD matrix into a 2D projection,[24] shown in Figure S3. We qualitatively confirmed the same results as in Figure 1: AF and OF are superimposed, while all the other models are well-separated.

In addition to clustering analysis reported in the main text, we also performed a structure-based sequence alignment of the cluster centroid structures shown in Figure 3, using the STAMP algorithm implemented in the MultiSeq tool[25] of VMD[26] (version 1.9.4a57). Such alignment (Figure S4) reveals that TM6 contains the largest number of mismatches within the helix, followed by TM7. This is in line with the differences in the TM6-TM7 interface described in the main text (see Figure 4).

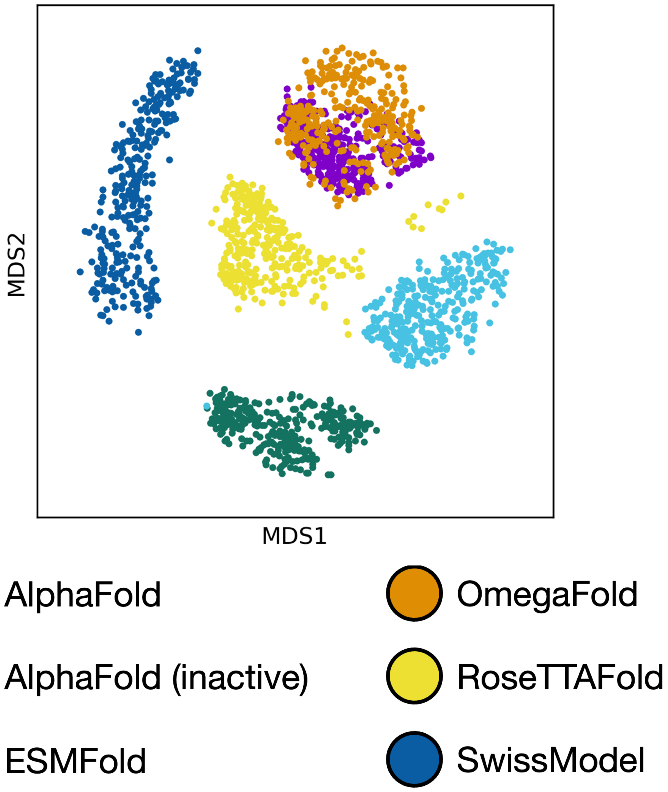


Figure S3: MDS-based similarity representation of all the sodium-bound hOR51E2 trajectories.

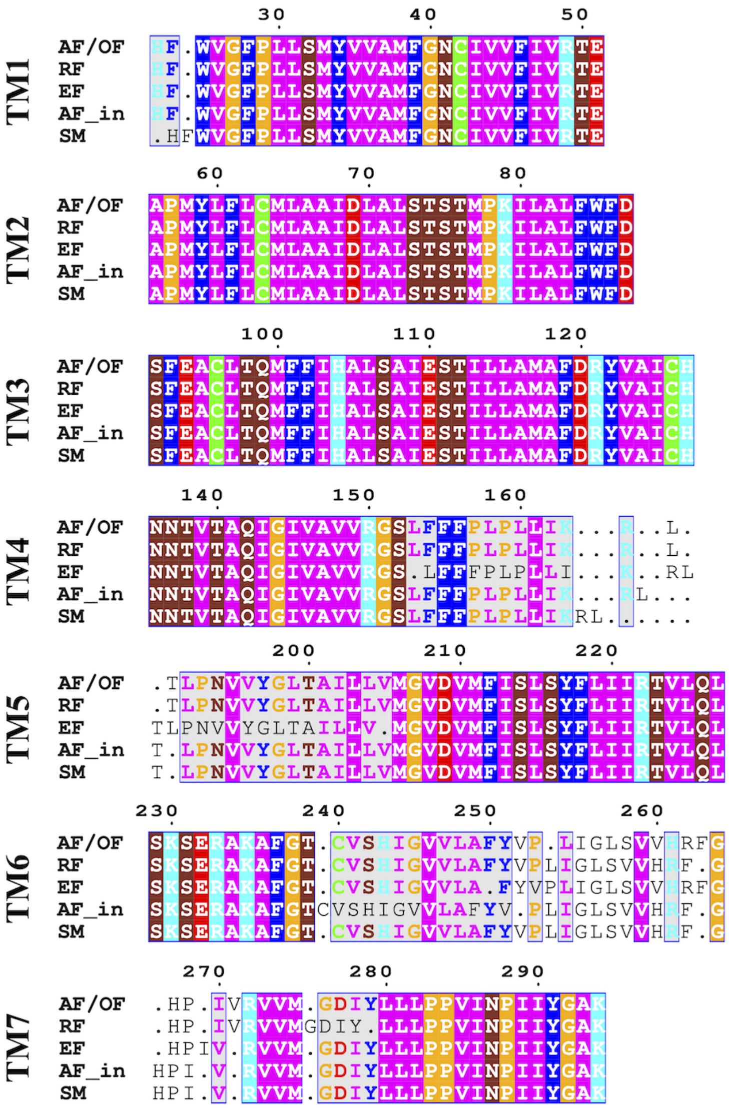

Figure S4: Structure-based multiple sequence alignment of the cluster centroid structures shown in Figure 3. The MSA was visualized using ESPript tool.[27] Residues identically aligned in all six models are displayed with white characters and enclosed in a box colored according to their physicochemical properties, residues showing a mismatch in one model are shown within a grey box and font color based on their properties, and gaps and residues mismatched in two or more models are shown in black font.

#### Comparison of the two best *in silico* models with the experimental structure of hOR51E2

We used the cryo-EM structure of hOR51E2 in the active state (PDB 8F76) to validate the two best inactive models according to our MD-based protocol, AF/OF and $AF_{in}$ (i.e. the centroid structures of clusters 1 and 4 in Figure 2). We first calculated the backbone RMSD of the seven transmembrane (TM) helical bundle, using the definition of the TM helices listed in Table S1 and the experimental structure as reference. The obtained RMSD values are 2.46 Å and 3.54 Å for the AF/OF and $AF_{in}$ models, respectively. However, such overall RMSD reflects both structural differences due to the quality of the models and conformational changes upon receptor activation. Indeed, the experimental structure was solved in the active state, whereas the computational models correspond to distinct inactive states (see Figure S1). Hence, we additionally quantified the structural changes due to the different activation state of the three structures using the 7x7 RMSD matrix of the individual TM helices, as defined in reference.[28] Such matrix was obtained with VMD[26] (version 1.9.4a57) by successively fitting one of the seven TM helices to the reference structure (here PDB 8F76) and calculating the backbone RMSD for each of the seven TM helices; for further details see the 7x7_rmsd.tcl script available in the zenodo repository (https://doi.org/10.5281/zenodo.7817679). The corresponding matrices are shown in Figure S5; each row represents the superposition to helix $i$ (S-TM$i$) and each column the backbone RMSD of helix $j$ of the model with respect to the same helix in the experimental structure (TM$j$:TM$j$). The TM helix showing the largest backbone RMSD is TM6, in line with conformational changes in this helix being necessary for activation of hOR51E2[29] in particular and class A GPCRs in general.[30] In this regard, it is worth noting that the AF/OF models were trained on both active and inactive GPCR experimental structures, whereas $AF_{in}$ only on inactive GPCR structures.

In addition, we analyzed (i) the residue-residue contacts at the interface between TM6 and TM7 and (ii) the side chain orientation of the residue lining the ligand binding site based on the cryo-EM structure. Such comparisons are reported in the main text and revealed a better qualitative agreement of the $AF_{in}$ model with the experimental structure.

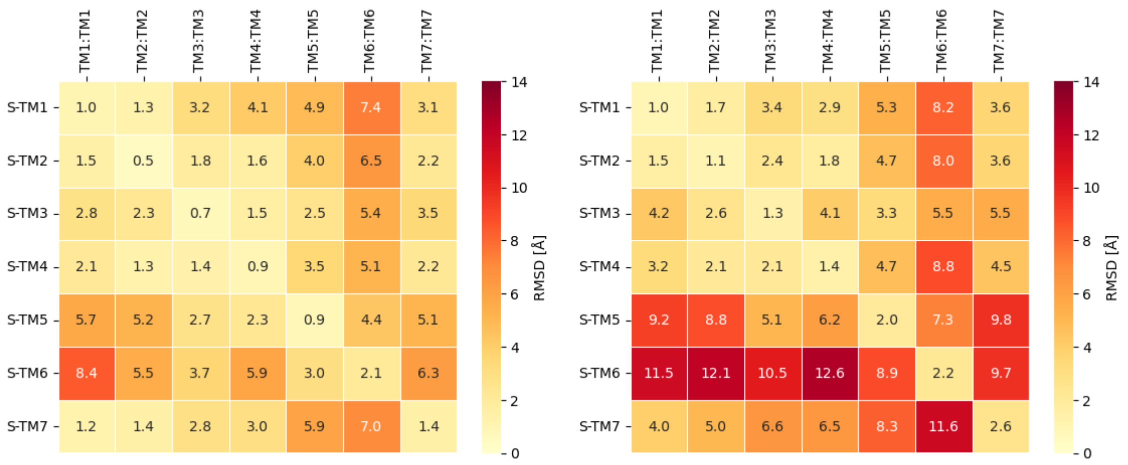


Figure S5: 7x7 RMSD calculation of AF/OF (Left) and $AF_{in}$ (Right) cluster centroids vs. the experimental hOR51E2 cryoEM structure (PDB 8F76).

Table S3: Conserved motifs in hORs with respect to class A GPCRs. Data taken from references[10,31–34]

| TM helix | hORs | class A GPCRs |
|---|---|---|
| TM1 | $G^{1.49}N^{1.50}xxI^{1.53}$ | $xN^{1.50}xxV^{1.53}$ |
| TM2 | $L^{2.46}S^{2.47}xxD^{2.50}$ | $L^{2.46}xxxD^{2.50}$ |
| TM3 | $D[E]^{3.39}$ | $S^{3.39}$ |
| | $M^{3.46}A^{3.47}Y[F]^{3.48}$ | – |
| | $D^{3.49}R^{3.50}Y^{3.51}$ | $D[E]^{3.49}R^{3.50}Y^{3.51}$ |
| TM4 | $W^{4.50}$ | $W^{4.50}$ |
| TM5 | – | $P^{5.50}$ |
| | $S^{5.57}Y^{5.58}$ | – |
| TM6 | $KAFSTCxSH^{6.40}$ | – |
| | $V[I]^{6.44}xxF[Y]^{6.47}Y[F]^{6.48}$ | $F^{6.44}xxxW^{6.48}$ |
| | $G^{6.49}x$ | $P^{6.50}$ |
| TM7 | $N^{7.49}$P7.50x$I[L]^{7.52}Y^{7.53}$ | $N^{7.49}P^{7.50}xxY^{7.53}$ |

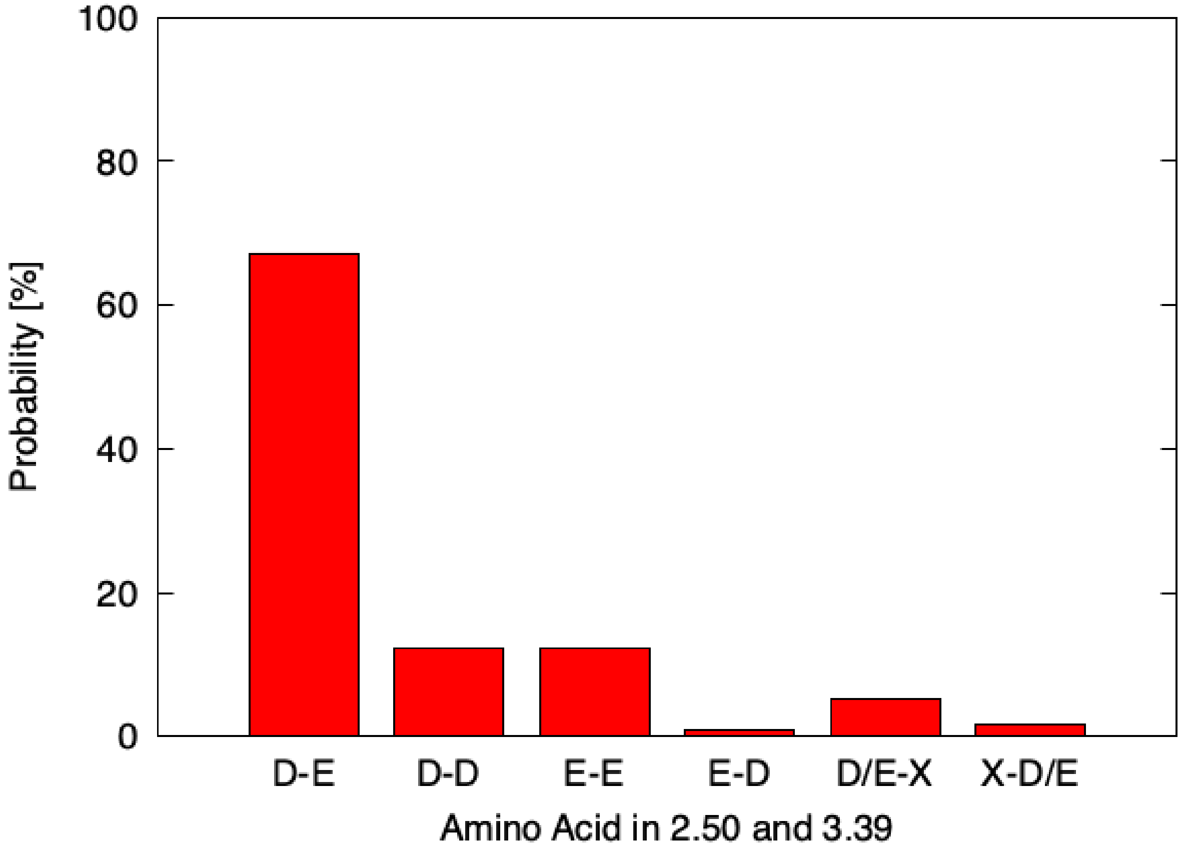


Figure S6: Residue conservation in BW positions 2.50 and 3.39 across human olfactory receptors, based on the MSA of 412 hORs in reference.[10]

Table S4: Residue conservation of the sodium pocket in hORs and class A GPCRs. The fifteen positions included were selected based on reference.[35] Data for hORs was taken from the MSA of 412 hORs in reference,[10] whereas for class A GPCRs it was aggregated from the 259 receptors analyzed in reference;[35] conservation frequency is indicated between parentheses as percentage. The two positions discussed in the main text (2.50 and 3.39) are highlighted in red.

| **Position** | **hORs** | **class A GPCRs** |
|---|---|---|
| 1.50 | N (99) | N (98) |
| 1.53 | I (62) | V (67) |
| 2.46 | L (93) | L (91) |
| 2.47 | S (74) / A (24) | A (75) |
| 2.49 | L (31) / A (19) | A (59) / S (24) |
| 2.50 | D (82) / E (16) | D (94) |
| 3.35 | F (43) / L (39) | S (20) / N (29) |
| 3.39 | E (81) / D (13) | S (76) |
| 3.43 | L (93) | L (76) |
| 6.44 | V (83) | F (79) |
| 6.48 | Y (68) / F (24) | W (67) / F (15) |
| 7.45 | T (50) / P (17) / I (15) | N (70) |
| 7.46 | P (96) | S (64) |
| 7.49 | N (98) | N (74) / D (20) |
| 7.50 | P (97) | P (94) |
| 7.53 | Y (96) | Y (91) |

(23) Bonomi, M.; Bussi, G.; Camilloni, C.; Tribello, G. A.; Banáš, P.; Barducci, A.; Bernetti, M.; Bolhuis, P. G.; Bottaro, S.; Branduardi, D.; Capelli, R.; Carloni, P.; Ceriotti, M.; Cesari, A.; Chen, H.; Chen, W.; Colizzi, F.; De, S.; De La Pierre, M.; Donadio, D.; Drobot, V.; Ensing, B.; Ferguson, A. L.; Filizola, M.; Fraser, J. S.; Fu, H.; Gasparotto, P.; Gervasio, F. L.; Giberti, F.; Gil-Ley, A.; Giorgino, T.; Heller, G. T.; Hocky, G. M.; Iannuzzi, M.; Invernizzi, M.; Jelfs, K. E.; Jussupow, A.; Kirilin, E.; Laio, A.; Limongelli, V.; Lindorff-Larsen, K.; Löhr, T.; Marinelli, F.; Martin-Samos, L.; Masetti, M.; Meyer, R.; Michaelides, A.; Molteni, C.; Morishita, T.; Nava, M.; Paissoni, C.; Papaleo, E.; Parrinello, M.; Pfaendtner, J.; Piaggi, P. M.; Piccini, G.;

Pietropaolo, A.; Pietrucci, F.; Pipolo, S.; Provasi, D.; Quigley, D.; Raiteri, P.; Raniolo, S.; Rydzewski, J.; Salvalaglio, M.; Sosso, G. C.; Spiwok, V.; Šponer, J.; Swenson, D. W. H.; Tiwary, P.; Valsson, O.; Vendruscolo, M.; Voth, G. A.; White, A. Promoting transparency and reproducibility in enhanced molecular simulations. *Nature Methods* **2019**, *16*, 670–673.

(24) *Modern Multidimensional Scaling*; Springer New York, 2005.
